# Simulation of Karyotype Evolution and Biodiversity in Asexual and Sexual Reproduction

**DOI:** 10.1101/481275

**Authors:** Andrew Y. Ying, Christine J. Ye, Hui Jiang, Steven D. Horne, Batoul Y. Abdallah, Guo Liu, Hao Ying, Henry H.Q. Heng

## Abstract

Whether sexual reproduction increases biodiversity remains controversial. Traditionally, sex within a species has been thought to increase genetic diversity, inferring an acceleration of macro-evolution, promoting biodiversity. Recently, it was suggested that the main function of sex is to maintain genome integrity, rather than increase genetic diversity or purify deleterious genes within populations, as the karyotype encodes/safeguards the genomic blueprint. As such, the contribution of sex to biodiversity needs to be re-examined. Since many simulation studies focus only on gene-level selection, it is important to investigate how sexual and asexual reproduction differentially impact patterns of genome-level evolution and biodiversity. Based on the key difference between sexual and asexual reproduction, that sexual individuals are required to mate with a partner of the same genome for successful reproduction, we have performed a simulation to illustrate how such differences impact genome-mediated biodiversity. Asexual populations displayed high genome-level diversity whereas sexual populations evidenced low genome-level diversity. Further analysis demonstrated that the requirement of finding a partner possessing a compatible genome prevents new sexual species from emerging, which may explain why geographic isolation can promote speciation: by increasing mating and survival-domination opportunities. This study challenges the traditional concepts of speciation and the function of sex.

## Introduction

Amid many hypotheses, evolutionary explanations for the main function of sex have often assumed that the adaptive advantage of meiosis arises from the genetic variants produced [1,2,3], which promotes population diversity and evolution [4-8]. For example, sex can create advantageous gene combinations by bringing together alleles of different individuals, and that sexual species can compete against parasites by constantly mixing genes to obtain novel, beneficial combinations and variants [9]. Other functions of sex that have been proposed include DNA repair and elimination of deleterious gene mutations in populations [10, 11]. Together, the gene mixing and/or purifying mechanism during meiosis explains the difference between sexual and asexual organisms (assuming asexual organisms possess nearly identical genomes). This viewpoint seems to fit well with current evolutionary concepts and serves as a key foundation for various popular theories as to why sex is advantageous to asexual reproduction, despite its heavy associated costs [9,12]. This concept is also inferred as the mechanistic link between micro- and macro-evolution, as the accumulation of small changes contributed by gene mixing is theorized to lead to emergence of new species over time.

Despite the popularity of this aforementioned framework, challenges remain:

first, each independent hypothesis appears valid on its own with certain evidence, but no one seems to be universally applicable. It is thus difficult to have a unified theory for the function of sex, despite the fact that almost every year, there are high profile publications from top scientific journals followed by popular press claiming that the mystery behind sexual reproduction’s existence has been finally solved. Second, some important paradoxes have been ignored (such as real distribution patterns of sexual and asexual species in harsh and drastically changed environments, the existence of self-sex, and the inconsistencies of many models that support the function of sex, especially the general assumption that asexual organisms display “identical genomes“) [12,13]. Third, under a fairly stable environment, there is no evolutionary incentive for individuals to undergo the risk of shuffling genes by meiotic recombination if current gene combinations fit well as the result of selection [3]. Fourth, and most importantly, the majority of modeling has been focused on recombination-mediated genetic diversity within species, and there are only very limited studies that directly address genetic diversity above the species level. Obviously, these concerns strongly suggest that the current paradigm may be missing something fundamental regarding the main function of sex.

An alternative theory surprisingly comes from cancer evolution studies, which reveal that genome stability serves as a major evolutionary constraint to karyotype maintenance [12, 13, 14, 15]. In agreement with the examination of the evolution of meiosis and self-sex [16, 17], as well as the constraints from genetic/epigenetic and ecological factors [18, 19, 20], it was proposed that the main function of sex is to serve as a filter at the genome level that eliminates any significantly altered genomes (karyotypes) and preserves the identity of the genome-defined system [12, 13, 14, 21]. This concept departs from the current paradigm of focusing on the evolutionary benefits of genes and gene combinations (including gene-level repair), and shifts the focus onto genome-level (karyotypic) changes in sexual and asexual populations. Moreover, if the karyotype represents a new type of genetic coding system for system inheritance [13, 21, 22. 23, 24], genome-level karyotype constraint becomes an essential means of passing the key genomic inheritance (blueprint) within a species—and therefore defines a species. It is thus reasonable to hypothesize that the chromosomal set or karyotype encodes system inheritance and significant karyotype changes directly contribute to macroevolution, including speciation, while gene mutations modify the existing genome-defined species [13, 14, 21]. Based on the above realizations, comparing how the function of sex contributes to the diversity of karyotypes and the biodiversity is of high significance.

One key prediction of our hypothesis is that sexual reproduction drastically reduces genome-level diversity, compared to asexual reproduction. Computer simulations may be useful in testing this prediction. In fact, such types of simulations are frequently used to compare the advantages of sexual and asexual populations under various conditions. However, most previous studies have been focused on the pattern of gene mutation and gene combination, population size and dynamics, and differential selection pressure, with often conflicted results [12,25,26, 27, 28]. One common basis has been the key assumption that sex generates diversity through meiosis while asexual reproduction produces identical “clones.” Though new sequencing data and conceptual syntheses suggest otherwise [12,13, 29], this assumption remains dominant. Recently, increased reports have analyzed the relationship between species diversity and gene-mediated genetic diversity [30,31]. Furthermore, sexual reproduction can paradoxically increase genetic variation but reduce species diversity [32]. These studies have forcefully illustrated that there is in fact no consensus regarding the important concept that observable population-level microevolution is sufficient to account for macroevolution for the large-scale changes over longer periods of life’s history [33].

By modeling genome-level changes (size [length] and configuration [topology]) in simulated sexual and asexual populations, we have analyzed the relationship between sexual and asexual reproduction, genome (karyotype) alteration, and resultant above-species-level biodiversity. The basic assumption of our simulation was that, unlike asexual reproduction, successful sexual reproduction requires an individual mate with a partner of the same genome. No assumptions were made regarding genetic diversity or advantages of either sex. The rest of the conditions are identical (identical rate of genome alteration, same types of genome (karyotype) alteration, same number of passages, same population size, and absence of selection factors). Following the simulation of 5000 generations, with constrained population size, genome-level diversity was found to be drastically higher in asexual populations, confirming our predictions. Since drastic genomic (karyotypic) alteration is directly linked to macroevolution, this simulation suggests that sexual reproduction may act to slow macroevolution. More importantly, however, this seemingly foreseeable simulation also revealed some big surprises: first, the number of new species of a visible population size formed from sexual reproduction was much lower than expected, with only the original species remaining dominant. Second, there was no gradual accumulation of biodiversity past generation 15: biodiversity levels remained the same quickly after an initial level of diversity was reached. Various parameters were incorporated into the simulations to represent the influence of factors such as rate of genome-level alteration, number of simulated generations, and the probability of mating among individuals with the same altered genomes as a result of population diversity. Together, these analyses provide new insight of how the requirement of mating defines the pattern of biodiversity, and support the hypothesis that geographical isolation might be needed to increase the probability of successful speciation from an established (ancestry) species, rather than serving as a precondition to stop the gene flow that generates genetic diversity within species.

In summary, based on the new finding that genetic diversity is lower in many sexual species than asexual ones, and the new realization that sex functions as a genome level constraint, we have focused on some more realistic and thus more relevant assumptions than many traditional ones (which either don’t fit the facts or represent secondary features of the function of sex). Further studies will include some traditional assumptions for additional comparisons.

## Materials and Methods

Our simulation study was based on four distinct stochastic models of fluctuations and alterations in populations through time, measured in generations. Specifically, these models were: (1) asexual reproduction with consistent rate of genome size changes (both increasing and decreasing), (2) sexual reproduction with consistent rate of genome size changes, (3) asexual reproduction with chromosomal inversions to represent topological genome changes, and (4) sexual reproduction with the same chromosomal inversions. Gene state mutations were also included in all models.

We modeled each individual of the simulated populations as a single-chromosome organism. Each chromosome was represented by a numerical string of length *N*, comprised of the numbers 1 to *N*, arranged in ascending order. Each numerical element of the string represented a gene and each gene had an associated numerical marker to represent a gene state. We assumed that each asexual individual was capable of self-reproducing an offspring, while each sexual individual required a partner of identical (in both length and topology) genome to reproduce. Each individual was able to survive and reproduce for five generations before “death” occurred. To simulate reality, a user-defined per-generation population cap was introduced to control population size.

The following are details regarding genome and gene manipulations for our simulations:

1. Genome size change model (represented genome-level variation): This model increased or decreased, with equal probabilities, the total number of genes in a chromosome by adding or removing genes from its end. We limited the range of alteration to 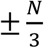 The probability of an alteration occurring each time an individual was created was denoted *α*_1_. The number of genes added or removed (*δ*_1_) in a given alteration was *θ*_1_ × *N*, where *θ*_1_ was a fractionizing constant between 0 and 1 (i.e. magnitude of gene change was *δ*_1_ = *θ*_1_ × *N*). Genomes of distinct lengths represented distinct karyotype-defined species (i.e., if chromosome length changes yielded chromosomes of lengths *N* −5, N, and *N* + 5, three species would be in existence). All numbers were rounded to the nearest whole number.
2. Gene order inversion (represented genome topology variation or re-organization of the preexisting genome): An inversion was the reversal of an ordered sequence of genes in a chromosome. More specifically, a gene within the *N*-length sequence was first randomly selected to be the point of inversion. This gene and the *δ* – 1 genes to the left of it then had their orders reversed (e.g., Genes 1, 2, 3, 4, 5 would be reversed to yield 5, 4, 3, 2, 1 if Gene 5 was deemed the point of inversion and δ = 5. Partial inversions were not allowed to occur (e.g., if Gene 4 were selected as the point of inversion and δ = 5, Genes 4 to 1 would not reverse as the full magnitude of inversion, 5, would be impossible to attain). The probability of an inversion occurring each time an individual was created was denoted *α*2. The number of genes reversed was calculated by taking *θ*_2_ × *N*, where *θ*_2_ was a fractionizing constant between 0 and 1, yielding a magnitude of gene change of *δ*_2_. As such, the range of points capable of being points of inversion for a *N*-length string was *δ*_2_ to *N* - *δ*_2_ + 1. It was also possible for subsequent inversions to occur within already-inverted sequences, thus allowing for many unique combinations. Chromosomes of distinct gene sequence orders (genomic topology or karyotype) represented distinct species (i.e., if gene sequence inversions yielded ten varying sequence orders, ten species would be in existence).
3. Gene state alterations (represented gene-level variation): This alteration was the change of one gene state to another in order to mimic allele changes. The probability of the event occurring each time an individual was created was denoted *µ*. In our four models, there were a total of ten gene states representing genetic diversity occurring at the gene level.

We assumed that genetic alteration did not result in infertility or any other complications that would impair reproduction (without any selective pressures). The two asexual models each began with one individual in the first generation. Every individual, including those with altered genome status (length or topology) based on designed rate of change produced one identical progeny per generation until death (individuals with new genome alterations were generated based on the designed rate within population). Each model ran for 5000 generations.

The two sexual models each began with two individuals in the first generation. To mimic the process of sexual reproduction, we designed and implemented a pairing process. This process represented mating between individuals with the same genome and prevented individuals from reproducing when they had differing genomes (by either size or topology). Only individuals with the same genome length (for the genome size alteration models) or of identical genome configuration representing sequence order within the genome (for the genome inversion models) were capable of pairing and yielding progeny. Pairing occurred by chance, with the probability of one individual finding a mate being 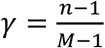 per attempt, where *M* was the total number of individuals alive, which was 125,000 after the generation population limit was reached (in fewer than 20 generations), and *n* was the total number of individuals of the same genome size or sequence order as the seeker. Note that the theoretical probability for the *k*-th seeker (for *k* < *n*) to find a mate was 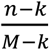 Hence, γ represented the first seeker’s probability (i.e., *k* = 1). Using it to approximately represent the probability for the rest of the seekers was reasonable because *M* was either much greater than *n* (for the less-represented genome sizes or sequences) or approximately equal to *n* (for the most dominant genome sizes or genome sequence configurations). This approximation simplified our model, enabling significantly reduced computing time. In our study, we allowed an individual to attempt pairing twice; consequently, the overall probability of a chromosome finding a mate was 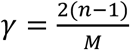 Every pair of matched individuals, including those with altered genomes, produced two replicas per generation until death. Table 1 shows other parameters of our four models utilized in conjunction with above-mentioned model parameters.

**Table 1.** Model Parameters. Four models tested with different parameters. *N* was the length of the numerical string representing a chromosome or genome. The initial number is 600 for Models 1 and 2, and 100 for Models 3 and 4. *α*_1_ and *α*_2_ denoted respectively the probability of the alteration and the inversion occurring each time an altered chromosome was created. *α*_1_ was for the genome size change model, *α*_2_ was for the gene order inversion (represented genome topology variation). θ1 and θ2 were fractionizing constants between 0 and 1 used in the genome size change model and the gene order inversion model, respectively. The probability of alteration occurring each time for gene state was created was denoted *µ*.

|  | **Sexual/Asexual** | **Genome**<br>**Alteration** | **N** | **$\alpha_1$** | **$\alpha_2$** | **$\theta_1$** | **$\theta_2$** | **$\mu$** |
| --- | --- | --- | --- | --- | --- | --- | --- | --- |
| **Model 1** | Asexual | Chromosome<br>length | 600 | 0.002 | N/A | 0.02 | N/A | 0.002 |
| **Model 2** | Sexual | Chromosome<br>length | 600 | 0.002 | N/A | 0.02 | N/A | 0.002 |
| **Model 3** | Asexual | Topology | 100 | N/A | 0.002 | N/A | 0.2 | 0.002 |
| **Model 4** | Sexual | Topology | 100 | N/A | 0.002 | N/A | 0.2 | 0.002 |

To execute our simulation study, we utilized MATLAB™, a widely-used engineering software package. The programs were optimized in terms of computing speed by taking full advantage of MATLAB™’s matrix manipulation commands and by streamlining program code, such as by minimizing use of FOR loops. A PC equipped with an Intel i7-2600 3.40 GHz CPU, 8 GB RAM, and 1 TB hard disk was used to run the MATLAB programs. Computing times for Models 1 and 3, which involved the genome length-change alterations, ranged from 4.9 hours (Model 1) to 11.5 hours (Model 3) when N = 600. The run-time of Model 4 was much longer due to the enormous quantity of gene-to-gene comparisons involved in the mating process. Computing times for Models 2 and 4 ranged from 11.4 to 120 hours, even when N = 100 and with program optimization. To keep computing time feasible, we curtailed the genome lengths in Models 2 and 4 to 100 and those in Models 1 and 3 to [400, 800] with the initial length being 600. It is important to note that using shorter genome lengths should not have compromised our results, as we experimented with varying genome lengths and found that the same trends manifested regardless of length used.

### Statistical analyses

We quantified and compared the two models in terms of biodiversity measured in two indices: richness (*R*, measured as the number of different variants), and the Shannon index (measured using the Shannon entropy 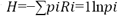, where *pi* is the proportion of individuals belong to variant *i*).

## Results

The straightforward simulations have generated the following results. These graphical figures are based on population data of the final five generations of individuals (generations 4996-5000). All figures describe model populations generated through the parameter values identified in Table 1.

### Asexual and sexual reproduction are linked to promotion and inhibition of genome-level variations, respectively

Figure 1 utilizes bar graphs to display the frequency of unique genome variations present in the final five generations of each simulation model. These results provide an idea of the overall effects asexual and sexual reproduction each have on the types of genome variation as well as the number of individual organisms over time. Evidently, as can be seen in Figs. 1a and 1b, asexual reproduction allowed for a wide range of species, represented by altered genomes (distinct genome lengths/unique gene sequence orders), to emerge consequent of lack of restriction. In the genome size alteration model, there were three apparent dominant species, plus seven visible ones. In the gene sequence inversion model, there were many more visible clusters (species), but each species had a low number of constituent individuals. Furthermore, even the non-dominant species were able to establish significant population sizes due to the lack of restrictions associated with asexual reproduction and, in Fig. 1b, were present in notable frequencies across the entire range of possible species types. Conversely, as can be seen in Figs. 1c and 1d, sexual reproduction acted as a strong restrictive force, filtering out virtually all large-scale genomic changes: in both the length change and gene sequence inversion models, only the original chromosomes remained dominant. Relative to the dominant species, the new species existed in infinitesimally small quantities due to the pairing process that mimicked sexual reproduction and inhibited mating among individuals with genomic variations. To allow for full appreciation of the disparities between species frequencies, it should be noted that the y-axis scales on the sexual models are much greater than those on the asexual ones.

**Fig. 1.**
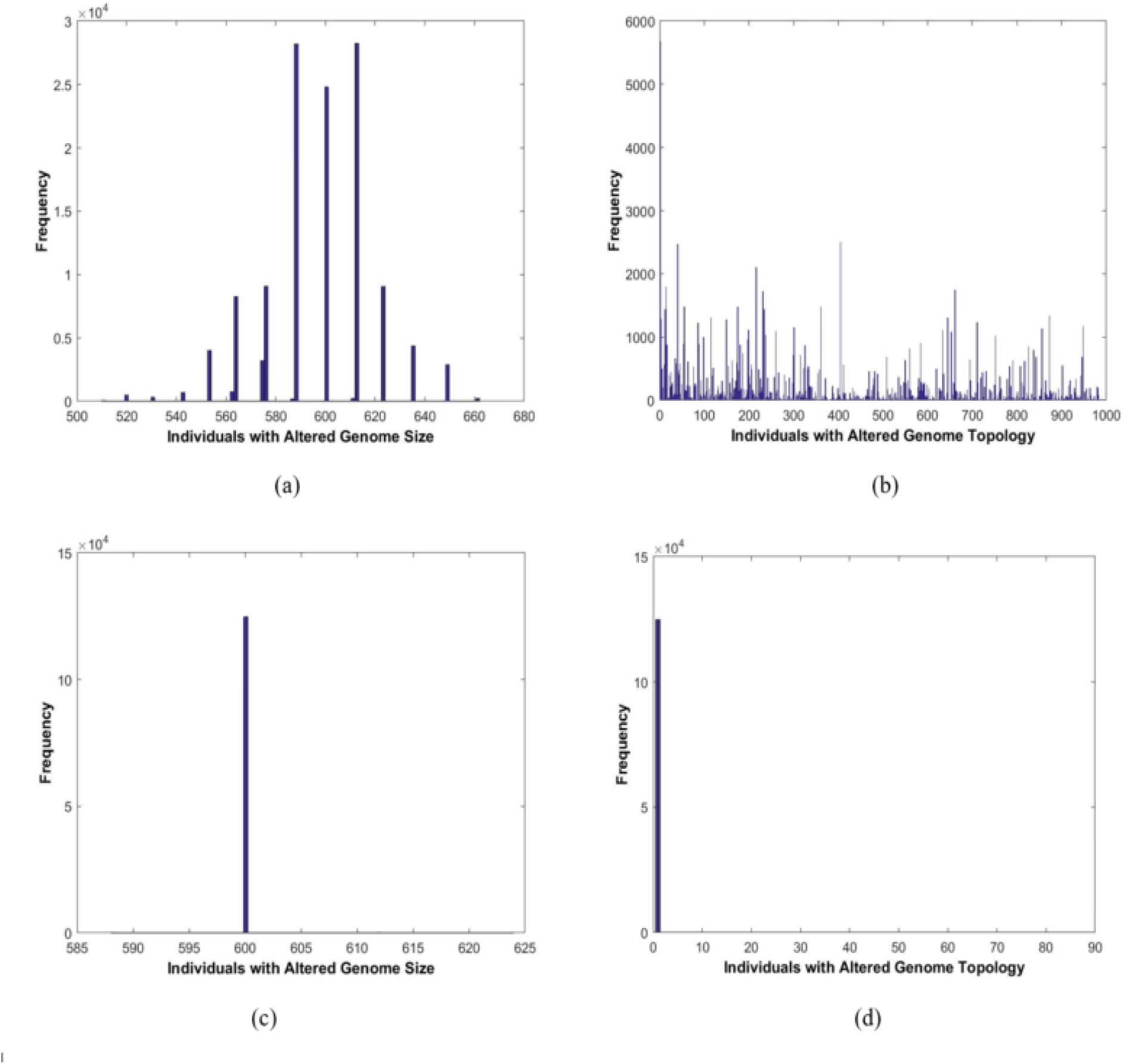
The frequency of unique chromosome variants: (a) Model 1, (b) Model 3, (c) Model 2, and (d) Model 4.

The five simulations for Fig 1a (Model 1) have richness indices of 26±2.9 (mean±standard deviation), and Shannon indices of 2.2±0.2. The five simulations for Fig 1c (Model 2) have richness indices of 3.2±0.4, and Shannon indices of 0.02±0.0. Using two-sample t-tests, the simulation results from the two models are statistically significantly different in terms of their biodiversity: p-value = 5e-5 for the t-test based on richness, and p-value = 1e-5 for the t-test based on Shannon index.

The three simulations for Fig 1b (Model 3) have richness indices of 1005 (mean±standard deviation), and Shannon indices of 5.50.08. The three simulations for Fig 1d (Model 4) have richness indices of 80, and Shannon indices of 0.020.0. Using two-sample *t*-tests, the simulation results from the two models are statistically significantly different in terms of their biodiversity: *p*-value = 5e-4 for the *t*-test based on richness, and *p*-value = 8e-5 for the *t*-test based on Shannon index.

For comparison, the statistical data of four models is listed in table 2.

**Table 2:** The statistical comparison of the four models in Fig 1.

|  | Model 1 | Model 2 | Model 3 | Model 4 |
| --- | --- | --- | --- | --- |
| Richness index | 262.9 | 3.20.4 | 1005 | 80 |
| p-value (t-test) | 5e-5 |  | 5e-4 |  |
| Shannon index | 2.20.2 | 0.020.0 | 5.50.08 | 0.020.0 |
| p-value (t-test) | 1e-5 |  | 8e-5 |  |

In conclusion, in addition to the distinctive patterns of visual distribution, the simulation results from the two models are statistically significantly different in terms of their biodiversity.

Even though the overall constraint of sexual reproduction is foreseeable according to our simulation assumption, the extremely high degree of constraint is rather surprising. We expected to see at least a few new dominant species emerge. If new species cannot be formed in our simulation, then there must be some fundamental parameters that have been ignored (this is further analyzed in “Increasing the chance of specific mating among individuals with the same altered genome can promote biodiversity of a sexual species“).

Figure 2 is comprised of colormaps depicting the range and frequency of unique species variants present in each model, as well as the extent of gene-level variation consequent of gene state mutations within each cluster of the individuals with the same genome and gene status. The graphs offer a different perspective, with the intensities of colors, or lack thereof, serving as visual indicators of genetic diversity yielded by each model. As Figs. 2a and 2b show, there is an unmistakably large amount of genetic variation present in the asexual models. In Fig. 2a, there are three obvious dominant species hotspots; however, the peripheral areas of the colormap still have noteworthy abundances of species. Like Fig. 1b, Fig. 2b exhibits near-absolute genetic diversity – nearly 1000 species existent out of a possible 1000 — supporting the notion that asexual reproduction, over thousands of generations, may result in countless divergences and deviations from an original species’ genetic identity. Starkly contrasting the results of the asexual models are Figs. 2c and 2d. In both, only a lone column of color is visible: all other combinations are either nonexistent or exist in such low frequencies that their color bars cannot be distinguished from the background. As evidenced by Figs. 2c and 2d, the original species are the only substantial populations even after 5000 generations have passed – this attests to the restrictiveness of sexual reproduction on genetic identity. Just as it was noted in the above discussion of Fig. 1, the scales of the colormap diagrams are drastically larger in the sexual models than in the asexual models. This result indicates that the mating requirement also impacts gene-level diversity.

**Fig. 2.**
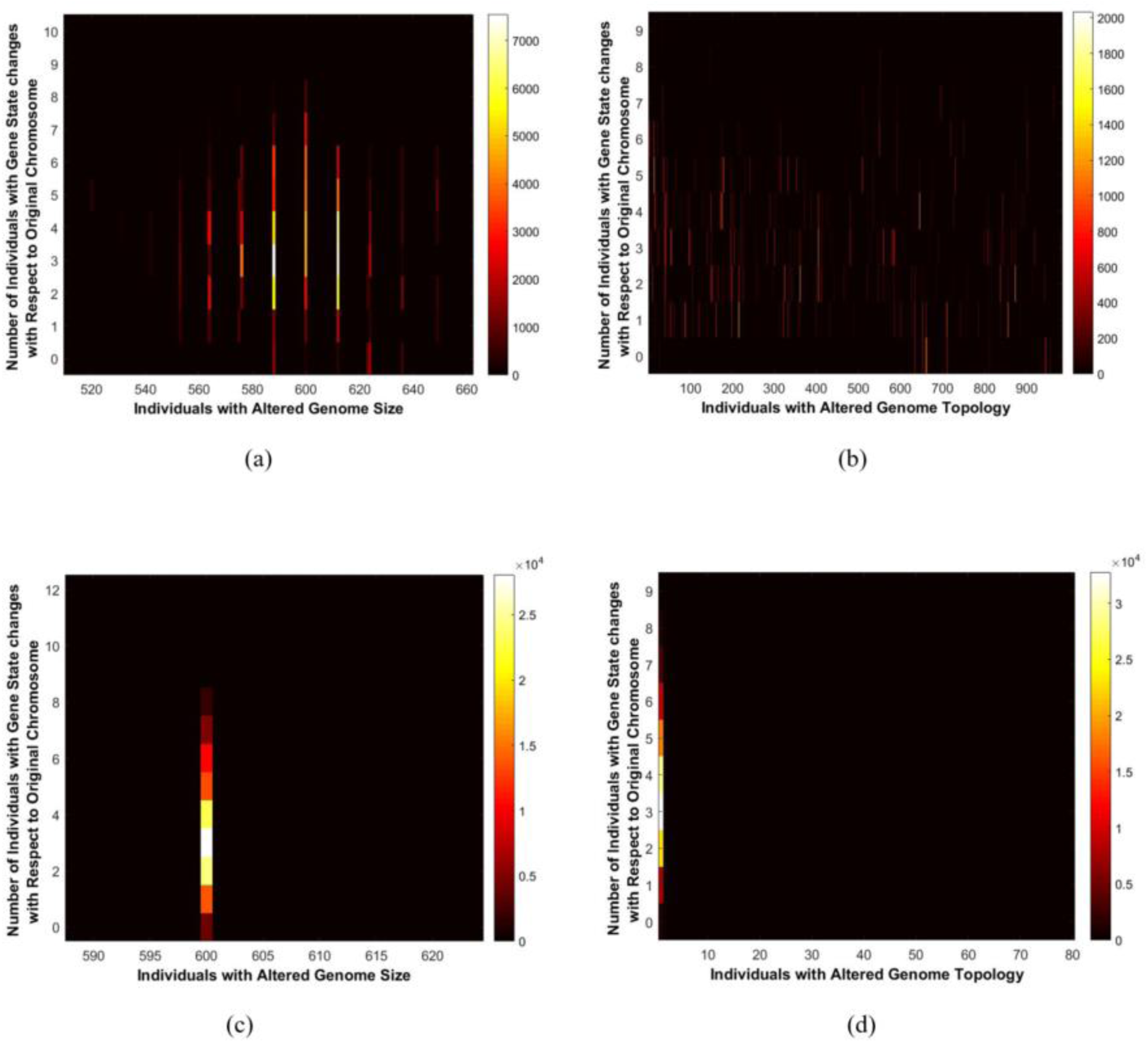
The range and frequency of unique species variants present in each model as well as the extent of gene-level variation consequent of gene state mutations within each species: (a) Model 1, (b) Model 2, (c) Model 3, and (d) Model 4.

### Rate of genome alteration contributes to genome-level diversity, especially in asexual reproduction

Based on the above simulations, it is worthwhile to change the rate of genome alteration, to investigate whether alteration rate plays a major role in genome-level diversity, as many have suggested. The rationale of using *α*(rate of genome change) = 0.002 is based on the fact that Down syndrome occurs in 1 of every 800 infants (0.0012). Since Down syndrome is the most common birth defect representing genome alteration, using *α* as 0.002 is reasonable and factors in other types of genome alterations. This rate is also comparable with rates other investigators have used. Additionally, the exact value of *α* is less crucial in our simulation, as both asexual and sexual reproduction used the same *α*. This point has been illustrated by model variations using *α* = 0.00002 and 0.2. We believe this range of *α* values covers that of reality.

Figure 3 displays via bar graphs the frequency of unique species variants (individuals with different genomes) in asexual and sexual genome size alteration models (Models 1 and 3) after the rates of genome size alteration have been increased or decreased 100-fold (compared to the rate used in Figs. 1a and 1c). In Figs. 3a and 3c, the rates of genome size alteration were decreased 100-fold. A remarkable decrease in the number of species was observed in the asexual population, while for the sexual population, there was no significant change when compared to Fig 1c. There is virtually no emergence of new species evident in Fig. 3c, the sexual reproduction model. In Figs. 3b and 3d, the genome alteration rate was increased 100-fold (compared to the rate used in Fig. 1a and 1c). A large increase in the number of emerged species occurred. For asexual Model 1 (Fig. 3b), the increased rate of genome alteration resulted in a high degree of genome-level diversity as evidenced by many individuals with all possible genome lengths being in existence in substantial quantity. However, although there was an increase in genome diversity in sexual Model 3 (Fig. 3d), there were still only five clusters of individuals or new species emergences, two of which were practically nonexistent due to miniscule population size. These rate changes of genome alteration and consequent differential diversity patterns of the populations further demonstrate the key difference between asexual and sexual reproduction in terms of producing genome-level diversity. Clearly, rate of genomic change affects degree of genome-level diversity, but much more dramatically in asexual populations than in sexual ones. In fact, in the sexual models, genome diversity only slightly increased even with high rate of genome alteration, indicating that sexual reproduction may be able to handle higher rates of genome alteration and maintain species genetic integrity. The related mechanism has been proposed as the “filter” to eliminate highly altered genomes [12]. Together with Fig. 1, the rate of genome alteration has been tested at 0.00002, 0.002, and 0.2. In all situations, the pattern of sexual and asexual reproduction-mediated genome diversity emerged highly different.

**Fig. 3.**
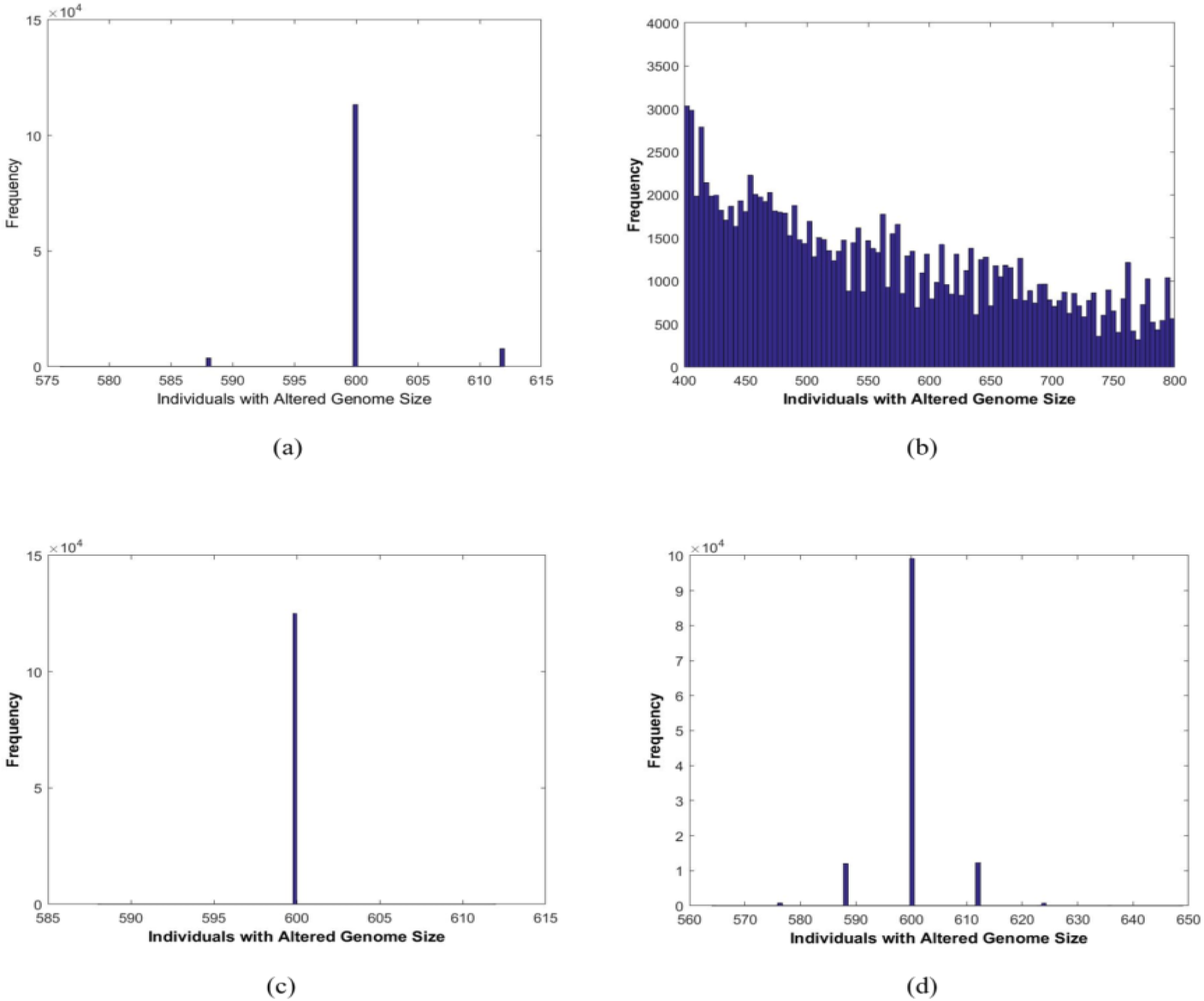
The frequency of unique species variants in each model after the rate of chromosome length mutations had been decreased or increased 100-fold: (a) Model 1 (rate (*α*) adjusted to 0.00002), (b) Model 1 (*α* adjusted to 0.2), (c) Model 2 (*α* adjusted to 0.00002), and (d) Model 2 (*α* adjusted to 0.2).

### Increasing the chance of specific mating among individuals with the same altered genome can promote biodiversity of a sexual species

The unexpected extremely high constraint of sex evidenced in Fig. 1 and 3 prompted us to identify the restrictive mechanism. Since there was no selective pressure in our simulation, such high constraint cannot be explained by differential selective advantages. If all new individuals with different karyotypes have the same fitness, what factor could prevent the relative dominance of a newly-formed species? In light of our random-mating model and the low rate of genome alteration in a large population, we hypothesize that new sexual species may be inhibited from becoming “visible” (achieving a certain number among all species) (Fig. 1) via lack of opportunity to mate with individuals of the same altered genomes. In other words, there are small numbers of newly-formed species in each generation, but due to the lower frequencies of successful pairing with each other (they have a much higher chance of meeting the original species), these individuals can only very rarely meet their own kind to reproduce successfully. These altered genomes will subsequently be diluted out of the population, and the original species will remain dominant. Contrastingly, in asexual populations, since there are no mating requirements, newly-formed genomes would be able to reproduce as successfully as the original when there is no selection.

We proceeded to hypothesize that, by increasing the chance of specific mating among individuals with the same types of altered genomes, a model parameter mimicking geographic isolation of new distinct species (e.g. on an island away from dominant parental populations) would enable more sexual species to become visible and form more dominant and thus visible clusters. To test this hypothesis, we introduce the condition to mimic geographical isolation that promotes the specific mating among individuals with the changed genome. For example, if we allow all individuals with altered genomes to find islands to separate themselves from the parental population, we essentially can simulate the hypothetical situation that every new, emerging species (with specific altered genome) is able to find its own distinct niche and isolate itself from all other pre-existing (parental) species. In this hypothetical situation, individuals would consequently experience a 100% success rate in meeting and matching with genetically-identical mates, as the individuals in each niche would share identical genomes. Fig. 4 presents with bar graphs the results of our sexual reproduction models with geographical isolation. Relative to baseline (no isolation factor) sexual model results depicted in Figure 1 (all individuals with newly formed genomes likely will mate with individuals of different genomes which will not lead to successful reproduction), a large increase in genome-level variation is observed. In the genome size alteration model, multiple species are able to establish considerable population size once given isolation, as opposed to one species emerging dominant as in the baseline sexual model. Although the original species, with N = 600, remains most dominant, sixteen distinct, partially-dominant species emerge (compared to just four in the baseline model). We observed similar results in the sexual gene sequence inversion model.

**Fig. 4.**
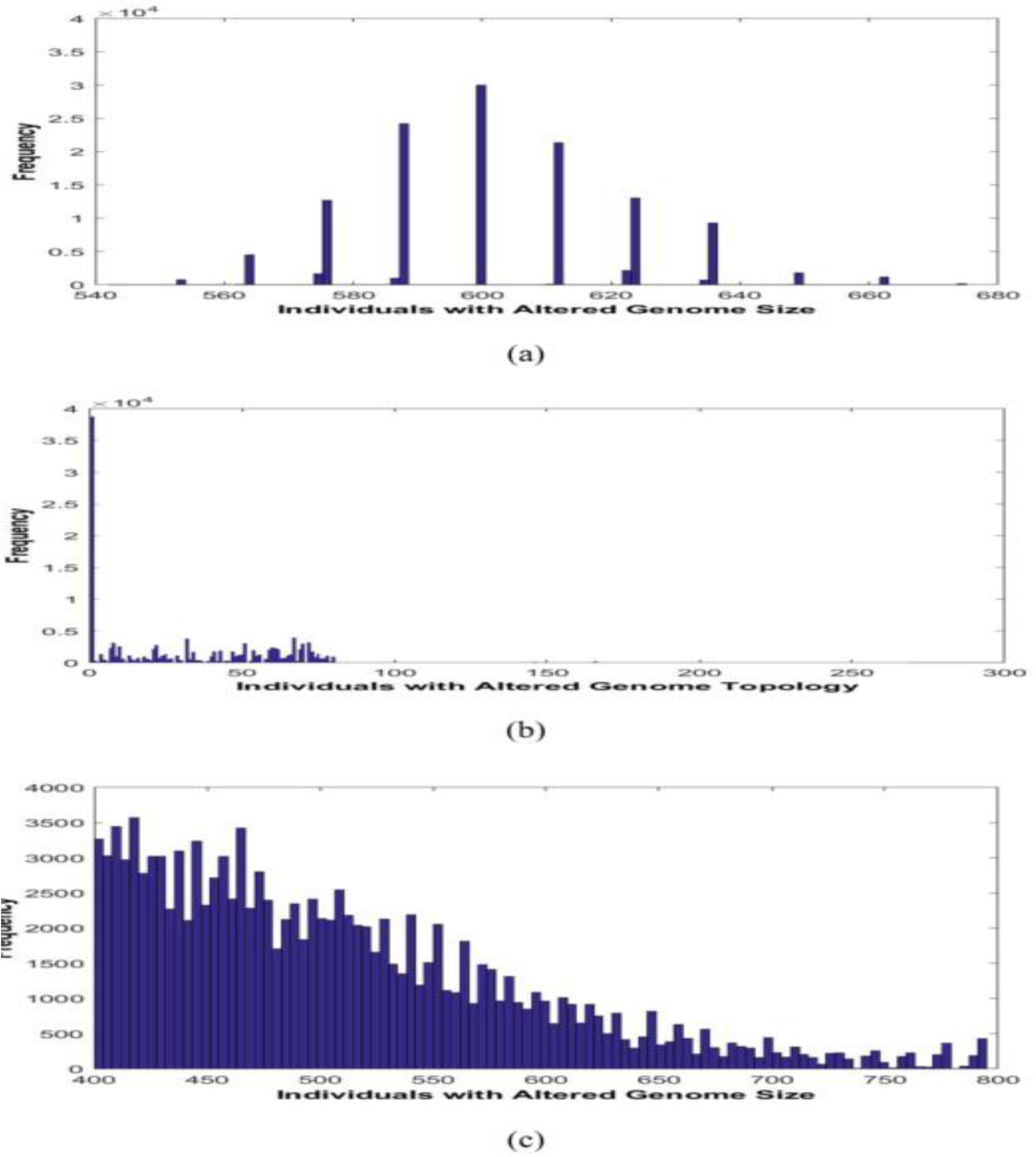
Hypothetical situation that in the sexual reproduction models every new, emerging species is able to find its own distinct niche and isolate itself from all other species, and individuals would consequently adopt a 100% success rate of matching with genetically-identical mates: Above: results after geographic isolation condition applied to: (a) Model 2, (b) Model 4, and (c) Model 2 (*α* increased to 0.2).

Although roughly the same number of species exists at the end of 5000 generations when compared to the baseline sexual inversion model, the population of each species has increased dramatically. Though the original species still has the largest population, four others are less than 300 members smaller: these results differ substantially from those of the baseline model, in which the original species had a population over 12000 members greater than that of any other. Our simulation seems to have provided an accurate and expected illustration of what would occur in sexual models containing an isolation factor. A more sophisticated simulation modeling selection would be able to further investigate and integrate fitness factors.

Recognizing the importance of isolation conditions in increasing likelihood of successful reproduction for new sexual species, we proceeded to investigate the impact of increased genome alteration rate in isolation. It is known that the number of emergent new species remains low when there is no isolation, even if genome alteration rate is increased (Fig. 3d). When alteration rate (α) was increased 100-fold to 0.2 with the isolation condition applied (Fig. 4c), large genetic diversity ensued (Fig. 4c). This suggests a powerful synergistic and compounding interaction between increased genome alteration rate and isolation conditions. A simulation for the genome sequence inversion model with *α* = 0.2 and the isolation condition was attempted, but could not be completed due to excessive computational demands.

### Overall degrees of genome diversity remain constant in sexual populations after 15 generations

To estimate how many generations of simulation would be needed in order to see regular patterns of genome diversity across generations, overall trend of genome diversity was compared using both the genome size alteration and topology alteration models. Such comparison would also reveal if there is an accumulative effect for degree of genome diversity over generations, as many believe there is accumulation of biodiversity over time. Fig. 5 exhibits through line graphs the total number of unique chromosome types, whether based on altered genome size or topology, in existence at each generation of the simulation for Models 2 (Fig. 5a) and 4 (Fig. 5b). As depicted by the graphs, near the start of the simulation, sharp increases in the number of unique genome types in existence occurred. However, this exponential increase did not last long and the total number of species in existence plateaued after about 15 generations at a limit of roughly 250 new species in both models. For the remainder of the simulation durations, the total number of species fluctuated around 250: in Models 2 and 4, for over 99% of the simulation’s operation time, the total number of unique genome types in existence was remained around 250, or 0.2% of the total population of 125,000 individuals. Because our genome size alteration rate was set to 0.2%, we would indeed expect the number of unique species to be roughly 250. Fig. 5 thus supports the technical integrity of our simulation by verifying we have simulated sufficient numbers of generations to yield statistically accurate results. In contrast, in our asexual populations, there was an accumulation effect: as the simulations progressed, individuals with altered genomes accumulated. Taken together, these results suggest sexual and asexual populations may behave differently to generate genome-level diversity. The constant low level of fluctuation might be caused by the genome constraint of sexual reproduction, and in terms of the population, the lower chance of specific mating. It is important to note that there was no long-term genome diversity accumulation effect in our sexual populations lacking isolation conditions.

**Fig. 5.**
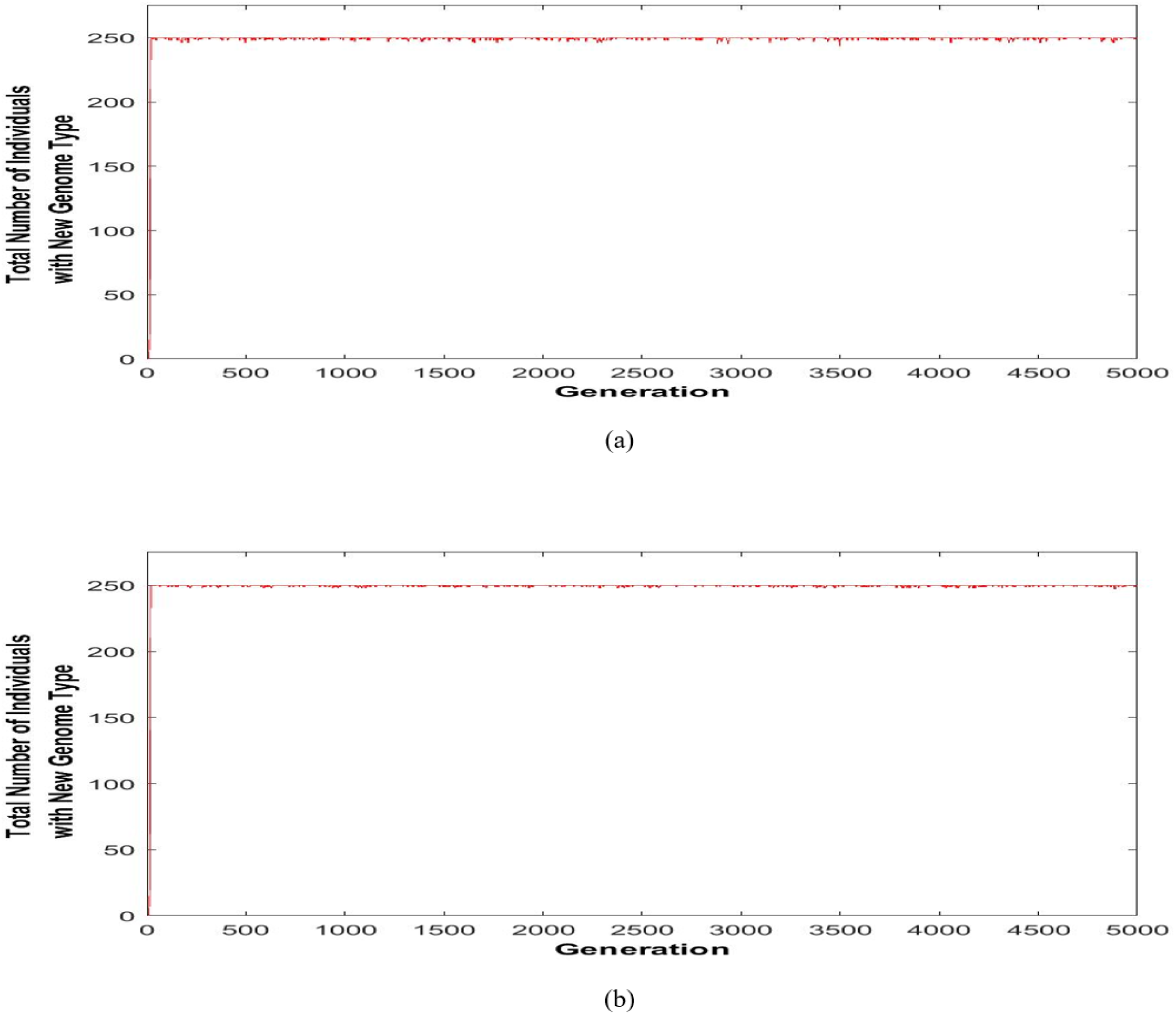
The total number of unique chromosome types in existence at each generation: (a) Model 2 (based on the number of distinct chromosome (genome) lengths), and (b) Model 4 (based on the number of sequence combinations (re-organized genome variations).

## Discussions

### The paradox of sexual reproduction: how does constraint promote biodiversity?

In discussion about the design of our simulation, one critical response has been that our results are unsurprising, as simulations are foreseeable (thus reducing its value). If sexual reproduction is modeled as requiring an individual find a partner with identical genome, it must of course reduce diversity, compared to an asexual model. However, our study surprisingly illustrates an obvious and profound paradox: if this simplest assumption is indeed true that sex requisites partnership and thereby reduces genome-level diversity as our simulation illustrated, why does the current paradigm hold that sex promotes biodiversity? Should we just challenge the paradigm based on this foreseeable concept/result? Furthermore, how could other factors, like meiotic recombination, DNA repair, and natural selection, achieve the transition from constraint to diversity? Clearly, this simple assumption and its consequence should be respected, even though it contradicts the generally accepted concept, because *it also fits the facts*. We hope our study may thus change the conversation, as the popular notion of mixed-gene adaptation fundamentally differs from the karyotype-constraint-related biodiversity herein described.

### Some considerations about key assumptions in our simulations

Like any other simulation, our study is both defined and limited by assumptions used. In many complex biological issues, when multiple factors are involved, there are often many assumptions which can serve as the basis for different simulations, and a specific simulation study can only reveal the logical implications of its assumptions. Traditionally, key assumptions on the issue of function of sex include increasing the genetic diversity through gene mixing by meiosis, accumulating genetic variations over time that lead to biodiversity, and geographic isolation promoting speciation by reducing gene flow. However, extensive decades-long research using these assumption-based models did not settle this important issue, leading to a paradoxical phenomenon: there are many solid individual simulation studies, but when considered all together, they do not make sense. Based on the new discoveries that the genomic diversity is much higher in asexual species than sexual ones, and that micro-cellular and macro-cellular evolution are fundamentally different, it is time to re-examine many previous assumptions to evaluate which are backed by facts, and which are the most real and fundamental. Three factors have emerged: 1) sex requires partners with the same or similar genomes, 2) genome variations can be simply achieved by changing the size and topology of the genome, and 3) the chance of individuals with “new” genomes mating with individuals of identical genomes is low when the vast majority of encountered individuals have incompatible “parental genomes“, especially when population size is limited and individuals do not have a large or unlimited number of mating opportunities. Many previous assumptions seem to be either incorrect (e.g. sequencing data does not support the assumption that sex increases genomic diversity) or secondary. We thus consider the main assumptions used in our simulation study as more realistic than many “traditional” ones.

These assumptions bear advantages and limitations. First, to simulate patterns of genome diversity based on a mating requirement is a novel approach. Despite knowing the assumption that the two parental genomes or chromosomes can pair if they have identical chromosome lengths and genomic topology could dictate the result that sex constrains variation, it was unknown what diversity patterns and trends would ultimately emerge from the simulation (how degrees of constraint and karyotype dynamics impact this process). In addition to supporting the function of sex as reducing genome diversity in general, our simulation reveals a striking signature pattern of continuous genomic alteration resulting from asexual reproduction. Prior to the simulation, we expected a fair number of unique-species clusters would emerge in our sexual populations. We did not expect that sex would be so restrictive that, under conditions where asexual reproduction would generate a large number of clusters, no dominant cluster emerged at all.

Second, it is well known that the reproductive barrier is mainly reflected at the genome level, as a majority of eukaryotic species have different karyotypes [14, 21, 23, 34, 35, 36, 37, 38, 39, 40]. Considering different altered genomes as different species in our simulation is thus reasonable. Of course, in natural biological systems there must be some degree of tolerance for differences in chromosome length or inversion state between two parental chromosomes. At this stage, we did not quantify the fuzzy level of tolerance, based on the fact that very few trisomies can be tolerated in humans, and that chromosomal inversion often leads to early spontaneous abortions. As for the observation that there are certain degrees of genome variation existing within each species [14,25], it is not known how many of these altered genomes actually contribute to reproduction. There must be a limitation towards the degree of allowable changes for a given karyotype, as the chromosomal number changes and inversions in humans are often related to unsuccessful reproduction. This in fact is the reason we have proposed the concept of the core genome of a given species [14]. One should consider individuals with altered genomes and their capacity, or lack thereof, to pass on altered genomes to offspring. Again, using humans as an example, despite the detection of trisomy 21 within populations, in most cases the extra copy of chromosome 21 cannot be passed down to new generations—even when both parents are trisomy 21. Creating or living with an altered genome is related to, but very different from, forming a new, viable species. According to genome theory, one should consider individuals with altered genomes and the capability of faithfully passing their altered genomes to the next generation as new species.

Third, this simulation is highly significant, as it links increased mating opportunity among individuals sharing identical genome alterations to genome-level diversity. It also explains why geographical isolation plays an important role for speciation, but in a non-traditional manner. Traditional explanations posit that natural barriers such as geographic separation of sub-populations, if sustained sufficiently long, may lead to the accumulation of independent genetic changes in groups and to mating incompatibilities [42,43]. A similar explanation is also used to understand topopatric speciation, a neutral speciation mechanism based on isolation by distance and assortative mating [30,44,45]. In contrast, in addition to the role of the sexual “filter” in preserving the “purity” of a species’ core genome [12,14,15], the connection between increased mating opportunity and visible emergence of new species represents a key factor to above-species-level biodiversity. Coupled with the observation that there is no long-term accumulated effect for sexual species to increase genome-level diversity over generations, geographical isolation can be seen to provide niches for newly-formed species to survive and become dominant (without direct competition or unsuccessful mating with ancestral populations), rather than simply providing an opportunity to accumulate genetic changes over time and eventually evolve into a new species (as the traditional view holds). This view is strongly supported by karyotype evolution in cancer [13,14,46]. The two phases (punctuated macro-cellular and stepwise micro-cellular) of evolution separate macro- and microevolution by a mechanism (genome vs. gene), rather than a time factor. Recently, clinical evidence has also supported this viewpoint. A recent cytogenetic analysis has revealed a stunning case of karyotype evolution in humans [47]. In a Chinese family, there was a Robertsonian translocation involving chromosome 14 and 15, generating a karyotype of 45 chromosomes (in the general population, the rate of Robertsonian translation is 1 in 1000). Due to mating between first cousins, a 44-chromosome homozygote karyotype was produced. Interestingly, when the 44-chromosome individual mated with a normal 46-chromosome individual, the resulting 45-chromosome child died. This case illustrated a few points: (1), karyotype changes can be achieved in as little as one generation. (2), a stable karyotype can be formed if mating can be achieved among individuals with the same heterozygotes. (3), to form a cluster of individuals of identical karyotype, mating must occur between individuals with the same karyotype. Otherwise, if the altered-genome individual mates with the original population, the altered karyotype can be eliminated. Evidently, for a new “species” of altered karyotype to establish a sizable population or become dominant, mating in isolation conditions is highly favorable.

Fourth, with the newly established platform, various parameters need to be modeled and tested in closer accordance to biological reality. For example, future experiments should be designed to test the impact of genetic recombination rate on genome-level diversity and determine the contribution of gene mutations to genome dynamics. With the assumptions used in our simulation, there is no direct relationship between gene- and genome-level alterations (Fig. 2). Such prediction has been supported by Figure 2. However, additional simulations are needed to include the accumulation factor at the gene level and its impact on the genome level, if any. In addition, only one chromosome is considered in our simulation. It is likely that this topic becomes much more complicated once multiple chromosomes and genetic recombination are modeled. Furthermore, the tolerance of genome alteration for mating needs to be simulated with biological data. Currently, there exist no quantitative data allowing for prediction of how many trisomies or chromosomal translocations successful sexual reproduction may tolerate. In our simulation, successful mating is highly simplified: we only required that two individuals with identical genomes “meet.” However, though there are many other factors that affect sex success rate, we believe they may not change the differences in genome-level diversity patterns/trends between sexual and asexual reproduction, as we tested a large range of genome alteration rates. Moreover, according to genome theory, these two levels are fundamentally different and possibly governed by different mechanisms (thus, the recombination data at gene and population level will not fundamentally change the biodiversity at the genome-defined above species level).

Fifth, as this simulation is mainly about genome-level (karyotype) diversity—which differs from copy number variations (CNVs), which are often not permitted by the meiotic pairing process—one should be careful to not confuse gene-level diversity with genome-level diversity. Genes (including CNVs) and genomes constitute fundamentally different levels of genetic organization and have differing evolution patterns [14,21]. Traditionally, chromosomes are considered the vehicles of genes, and karyotype-mediated speciation is overlooked [48]. Now, since the karyotype defines system inheritance and the identity of a species, the importance of the karyotype in speciation needs re-evaluation [22,23,49,50]. From the gene point of view, for example, inversions are often discussed in the context of local adaptation and linkage effects. The emphasis is on gene-level selection and gene frequency within a given population [51,52]. From the genome point of view, the order change within a genome represents a system coding change, and is an emergent property of the entire genome: the emphasis is on macro-evolution (more discussion can be found from the genome theory of cancer evolution, see [13,14]. Another example is selection type. Since there is no selection involved in our simulation, our results should not be explained as a particular form of purifying selection. Whereas purifying selection always reduces deleterious diversity. In our case, however, we only monitored overall biodiversity—without distinguishing beneficial or deleterious alterations.

### Significance of comparing genome-level diversity between sexual and asexual reproduction

How sex leads to the genetic variation that accelerates speciation has been debated for over a century. There are ample amounts of conflicting experimental and theoretical evidence and the newly-realized challenge is to separate genetic diversity from genome diversity [12,15,53,54]. In a recent theoretical study, it was shown that sexual reproduction could increase genetic variation but reduce species diversity [43]. Our simulation has not only supported their conclusion, but also offers explanation behind their once-seemingly paradoxical results. The answer may lie in the different contributions of gene- and genome-level change to micro- and macro-evolution, respectively [13,14,21]. On one hand, it is well known that gene mixing through meiosis can increase gene-level genetic diversity, despite that the initial function of meiosis is not to promote genetic diversity [17]. On the other hand, after demonstrating that sexual reproduction drastically reduces genome-level genetic diversity, we can now reconcile that sex can increase gene-level variation while reducing species (genome-level) diversity, and that a high microevolutionary rate may not necessarily yield a large number of species in natural ecosystems (macroevolution). To see sex as having dual roles may explain the key paradox involving active short-term evolutionary adaptation (microevolution) and long-term system stasis [15,55,56], even though the role of gene mixing might be much smaller than previously thought based on the concept of fuzzy inheritance [14,21].

It is now the time to appreciate the key differences between sexual and asexual reproduction, which require different evolutionary mechanisms, in evolution. Using somatic cell evolution as a model, it has been demonstrated that there are two phases of cancer cell evolution: a stepwise Darwinian evolution phase where gene mutation and epigenetic regulation dominate, and a punctuated macroevolution phase where genome replacement dominates. The phase transition is defined by overall system stability [14,21,46,50]. Though cancer cells can evolve through both gene and genome changes due to lack of a sexual filter, genome changes are the primary driving force for new systems that emerge with new genomes. In contrast, for the evolution of sexual organisms, sex functions as a system-stabilizing force that preserves system inheritance and constrains genome-level diversity, which supersedes gene-level diversity when in the context of new system formation. While sex permits and can sometimes even promote gene-level dynamics, it restricts significantly-altered genomes from remaining in the genome pool of a given species. This realization suggests that the evolutionary pattern of asexual reproduction fundamentally differs from that of sexual reproduction. Specifically, sexual organisms may accumulate genetic change during both micro- and macroevolution, whereas sexual organisms may only do so in microevolution. When macroevolution occurs in sexual organisms, new species emerge. Yet, whether the newly-formed species can become visible (by accumulating certain numbers) or dominate may largely be due to chance. As a result, for most asexual species, there exist high degrees of continuous diversity among them (a gradient). This has been recently supported by the sequencing of large numbers of ocean micro-organisms, and by the fact that it is quite difficult to identify their consensus sequences [48]. In contrast, for sexual organisms, there exist well-defined core genomes [13,23,34,38,56,57,58,59]. Despite high gene-level variations, the genomes of sexual organisms remain, for the vast majority of individuals in a given population, largely the same across generations.

In summary, we present a simulation based on genome-level changes (size and topology) to analyze the relationship between sexual and asexual reproduction, genome alteration, and resulting biodiversity. Our simulation data support the hypothesis that sex reduces genome-level (karyotype) diversity and slows macro-evolution. Based on this hypothesis, it appears less likely that the accumulation of gene-level diversity by meiosis will, somehow, lead to macro-evolution. This conclusion is in contrast with the popular view that genetic recombination increases gene and population diversity, as well as diversity above the species level, as population diversity is a necessary contributing factor to speciation.

While it is obvious that meiosis can increase genetic diversity at the gene level as gene combinations are constantly changing, we posit that the most important role of meiosis is actually genomic preservation via elimination of significantly-altered genomes and reduction of the chance of successful mating between individuals sharing altered genomes. These perspectives have not previously been vigorously simulated. According to genome theory, a systematic understanding of the main function of sex requires separation of the different types and degrees of genetic diversity that occur at the gene- and genome-level. While gene-level diversity is essential for microevolution, genome-level diversity is essential for macroevolution and biodiversity above a species level. The potential dual function of sex needs to be systematically studied. The idea that sex reduces biodiversity can be divided into two major mechanisms: (1) within a population of a given species, sex filters the passage of significantly-altered genomes; (2) when individuals with altered genomes are produced, the likelihood of new species formation is further reduced by stringent mating partner requirements imposed by sex that are exacerbated by lack of isolation conditions. By illustrating these mechanisms, our simulation demonstrates why it may be so difficult for speciation to be observed in sexual organisms, despite the occasional appearance of altered genomes. However, when the environment can produce extremely high levels of stress, genome instability can lead to genome chaos, the process of rapidly generating massive karyotype changes [60]. Under such conditions, selective pressure is so high that it will wipe out the majority of the original species, and only the newly emergent species have a better chance to meet and populate successfully. This simulation thus has illustrated a partial mechanism of this process. Clearly, these concepts depart from the neo-Darwinian viewpoint of how speciation occurs; thus, the issue requires immediate research for its further development and validation.

## Acknowledgements

This manuscript is part of a series of studies entitled “The mechanism of somatic and organismal evolution.” We would like to thank Drs. Root Gorelick and Yong-Bi Fu for their suggestions.

Author Contributions
Ying AY, Ying H and Heng HH designed experiments, Ying AY and Ying H performed the simulations; Ying AY and Heng HH drafted the manuscript; Hui Jiang performed the statistical analyses; Horne SD, Abdallah BY, Ye CJ and Liu G anticipated the discussion, literature search and editing the manuscript.

